# The Interplay of Prior Information and Motion Cues in Resolving Visual Ambiguity in Agent Perception

**DOI:** 10.1101/2024.05.30.595378

**Authors:** Sena Er, Hüseyin O. Elmas, Burcu A. Urgen

**Author notes:** Corresponding author *Email address:* (Sena Er). Authors equally contributed to this work (Sena Er, Hüseyin O. Elmas).

## Abstract

Agent perception is essential for social interaction, allowing individuals to interpret and respond to the actions of others within dynamic environments. In this study, we examined on how prior knowledge and motion cues are integrated to influence the temporal dynamics of perceiving agents. In order to make realistic but ambiguous stimuli in motion and form characteristics, we used human, robot, and android agents. Using temporal representational similarity analysis (RSA) on EEG recordings, we analyzed the representation of agent identities under varying conditions—Still versus Moving stimuli and Prior versus Naive contexts. Our findings revealed that prior knowledge and motion cues interact to produce distinct temporal patterns of representation. In the naive condition, information about the agent persisted longer during still presentations than during moving ones, suggesting that the processing of agents depends on the availability of motion information and prior information. Moreover, motion information affects the temporal processing of agents when no prior information about agents is available. These results highlight the critical roles of both bottom-up sensory inputs and top-down expectations and their interactions in resolving the ambiguities inherent in agent perception.

## 1. Introduction

Agent perception, the cognitive process of interpreting others’ actions to navigate complex, dynamic environments, underpins essential aspects of human interaction and decision-making. This fundamental cognitive ability enables us to discern intentions, assess threats, and respond to human and non-human agents by integrating motion cues and contextual information. Embedded in our evolutionary heritage, agent perception is not only vital for survival but also shapes social interactions and behaviors. However, the perception of agents is complex and ambiguous due to environmental noise and the fleeting nature of actions, necessitating a sophisticated interplay of bottom-up and top-down processes.

In elucidating the bottom-up processing of agents, the collaborative roles of the ventral (‘what’) and dorsal (‘where/how’) visual pathways are key, integrating body form and motion cues to form a coherent understanding of an agent’s actions and identity (Mishkin et al., 1983). This process aligns with theoretical models that propose distinct mechanisms for processing these cues. On the one hand, Lange and Lappe (2006) suggests a sequence-based recognition of movements through static postures. On the other hand, Giese and Poggio (2003) advocates for a parallel feedforward processing model, which posits that form and motion are analyzed independently within the ventral and dorsal streams, respectively. These models might not fully capture the complexity of agent perception. For instance, ‘Motion-blind’ patients can still recognize biological motion despite dorsal pathway impairments, suggesting alternative processing routes (Vaina et al., 1990; McLeod, 1996). These may involve cross-talk with the ventral stream (Beintema and Lappe, 2002; Jellema and Perrett, 2003; Gilaie-Dotan et al., 2013) or top-down influence from the dorsal pathway (Bar et al., 2006; Markov et al., 2013), challenging the model’s emphasis on distinct feedforward processing. Further expanding the dual-pathway model, recent findings highlight a third visual pathway dedicated to social perception, originating in the primary visual cortex and extending to the posterior superior temporal sulcus (Pitcher and Ungerleider, 2021). This pathway is particularly tailored for processing dynamic social stimuli, such as moving faces and bodies. It plays a pivotal role in social cognition and emphasizes the significance of motion in agent perception.

Action observation research further illuminates the mechanisms behind recognizing and interpreting others’ movements and intentions. The Action Observation Network (AON), spanning premotor, parietal, and occipitotemporal regions, is pivotal in mediating processes behind processing agents (Caspers et al., 2010; Nelissen et al., 2011). The posterior middle temporal gyrus and posterior superior temporal gyrus, part of the occipitotemporal node, are responsible for understanding agents and the social cues present in actions. Additional studies support the idea that the occipitotemporal cortex (OTC) is critical in processing the social aspects of observed actions, including agents (Orban, 2018; Sliwa and Freiwald, 2017; Tarhan and Konkle, 2020; Wurm et al., 2016). This interplay of neural pathways underscores the broader neural mechanisms, such as recurrent processing and the influence of prior knowledge, which are crucial for interpreting ambiguous visual stimuli. Recent studies point to the importance of recurrent processing for resolving visual ambiguities (O’Reilly et al., 2013; Wyatte et al., 2014; Kar et al., 2019; van Bergen and Kriegeskorte, 2020; Kornmeier et al., 2021), and the impact of top-down influences of prior expectation on biological motion perception, action observation, social perception (Hunt and Halper, 2008; Ondobaka et al., 2014; Schenke et al., 2016; Jacquet et al., 2016; Bach and Schenke, 2017; Chambon et al., 2017). Further supporting the necessity of recurrent processing, Kietzmann et al. (2019) demonstrated through magnetoencephalography (MEG) and deep neural network modeling that recurrence is crucial within the human visual system. Their research revealed an intricate dance of feedforward and feedback influences across the ventral visual stream. This supports the notion that agent perception involves both the emergence of low-level visual features in early areas and the modulation of these features by high-level categorical information in a top-down manner. Furthermore, biological motion perception, integral to agent recognition, also benefits from prior information, especially under ambiguous conditions (Hunt and Halper, 2008). This reliance on prior knowledge emphasizes the insufficiency of bottom-up processing alone for understanding complex visual social stimuli.

Overall, these insights necessitate reevaluating our understanding of how the brain processes visual social stimuli. They reveal a complex interplay of mechanisms enabling the brain to navigate interactions with others by utilizing top-down and bottom-up information. This involves intricate lateral, recurrent, and top-down communication between various brain regions. Synthesizing these top-down and bottom-up processes is essential for managing the perceptual ambiguity often found in agent perception. This dynamic suggests that the brain integrates low-level sensory data with top-down predictions to create coherent representations of agents, emphasizing the need for comprehensive models that accurately capture the complexity and ambiguity of agent perception in natural settings.

### 1.1. Non-Biological Agents as Ambiguous Stimuli

The investigation of visual perception frequently employs ambiguous stimuli to thoroughly explore the dynamics between bottom-up and top-down processing (Sterzer et al., 2009). Examples of such stimuli include luminance or contrast modulations (Pantle and Turano, 1992; Salge et al., 2020), the introduction of Gaussian noise (Salge et al., 2020), Mooney images Flounders et al. (2019), Necker cubes (Kornmeier et al., 2021; Hardstone et al., 2021), and a variety of optical illusions (Hardstone et al., 2021; Bill et al., 2022). These tools are pivotal in the literature for generating conditions under which visual cues can be interpreted in multiple ways. By doing so, they necessitate a greater reliance on top-down processes for resolving perceptual ambiguities. However, introducing ambiguity in dynamic and social stimuli through methods like dynamic noise addition (as in (Neri et al., 1998; Platonov and Orban, 2016)) or brief exposure durations (as in (Thurman and Grossman, 2008; Thurman et al., 2010)) involves a trade-off between manipulating sensory evidence and preserving ecological validity.

Exploring ambiguity in the context of agent perception has led to innovative approaches, such as employing non-biological agents like robots (Saygin et al., 2011; Urgen et al., 2018). Recent research indicates that non-biological agents activate AON similar to biological agents (Saygin et al., 2011; Gazzola et al., 2007). Saygin et al. (2011) found that human and robot agents similarly activated all AON regions except for the EBA (extrastriate body area), which exhibited more suppression for human-like appearance; suggesting that except for EBA, AON is not selective for the appearance or motion of agents. Interestingly, the bilateral aIPS (anterior intraparietal sulcus, a key node in the action observation network) showed similar responses for human and robot agents but demonstrated elevated suppression for Android agent, possibly due to prediction error caused by form-motion incongruity in line with the findings of (Cross et al., 2011; Kupferberg et al., 2017).

The ability of non-biological agents to evoke similar neural and cognitive responses to biological agents while inducing different activations based on incongruences in motion and form characteristics makes them powerful tools for investigating agent perception in ambiguous contexts.

### 1.2. Current Study

Despite recent advancements, the existing research on agent perception reveals considerable gaps, especially in understanding how low-level cues like motion and form interact with top-down influences such as prior knowledge of agents. The nuanced dynamics of how these factors collectively impact the temporal unfolding of agent perception remain largely unexplored.

Our study seeks to bridge this gap by examining the effects of prior knowledge and motion cues on the temporal dynamics of agent perception within the human brain. We focused on three distinct agents—**Human**, **Android**, and **Robot**—engaged in a variety of actions. The **Android** and **Robot** agents share the same underlying machinery, with the **Android** designed to mimic the **Human**’s appearance. Consequently, the **Human** agent embodies both biological appearance and movement; the **Robot** agent represents mechanical appearance and movement; and the **Android** agent merges a biological appearance with mechanical movement, mirroring the **Human**’s looks and the **Robot**’s movements.

We presented stimuli in video and static image formats to manipulate motion information. We also varied participants’ prior knowledge about the agents through two separate experiments: one where participants were pre-informed about the agents’ identities (**Prior Experiment**) and another where they had no prior information (**Naive Experiment**). This design enabled us to investigate the role of top-down processing when the form and motion cues provide ambiguous signals about agent identities. Our analysis aimed at uncovering how prior knowledge and motion cues influence the temporal dynamics of agent perception. We analyzed the temporal dynamics of agent perception using temporal RSA with multiple regression to understand how much of the information in the EEG signal is explained by the agent information while regressing out potential confounding factors such as luminance and optic flow.

We hypothesize that the temporal dynamics of agent representation are dependent on a combination of motion cues and prior knowledge. When both cues are present, they interact with each other to create a more nuanced perception of agent identity over time. However, when motion cues are absent, the effect of prior knowledge becomes even more pronounced, possibly because the lack of motion cues increases the ambiguity in agent identities, making the effect of prior knowledge even more significant.

## 2. Methods and Materials

### 2.1. Participants

The study involved 35 individuals from the UC San Diego student body, with 20 participating in the **Prior Experiment** and 15 in the **Naive Experiment**. All participants were free from neurological disorders and had normal or corrected-to-normal vision. They received course credit for their participation. Approval for the study was granted by the Human Subjects Ethics Committee at UC San Diego. Data for Experiment 1 were sourced from the work of Urgen et al. (2018).

### 2.2. Stimuli

The research employed a mix of short videos (lasting 2 seconds, termed as **Moving** stimuli) and static images (**Still** stimuli), showcasing three types of agents: **Human**, **Android**, and **Robot**, engaged in 8 actions. As **Android** and the **Robot** share the same underlying machinery, they have identical movements but distinct forms. Designed after the **Human**, the **Android** visually aligns with the **Human**. This design intricacy makes it harder to differentiate between these agents using only form or motion alone. The stimuli were displayed as either videos or images to manipulate the presence of motion cues. This method investigated how agent recognition occurs when their visual characteristics are either intact or limited. In the video segments, all agent types were depicted performing eight specific actions: drinking, hand examining, waving, body turning, table wiping, nudging, self-introducing, and paper throwing. **Still** stimuli were obtained from the initial frames of the respective **Moving** stimuli. Additional information on the stimuli is available in Urgen et al. (2018).

### 2.3. Experiment Design and Procedure

This study was structured around two EEG experiments differentiated mainly by the prior knowledge provided to participants. In the **Prior Experiment**, participants were introduced to the agents through a preliminary briefing. On the other hand, in the **Naive Experiment**, participants were not introduced to the stimuli set. They were only informed that they would view images and videos and provide judgments. Both experiments manipulated the agent’s identity (**Human**, **Android**, **Robot**) and the availability of motion cues (**Moving** or **Still**).

### 2.4. Prior Experiment

Participants in the **Prior Experiment** viewed videos in a pseudo-random order to avoid immediate repetitions. Videos were either presented alone or followed by their corresponding static image (the video’s first frame) for a random duration of 600–1000 ms. Before each video or image, a fixation cross appeared on the screen for 900–1200 ms, randomized for timing. Engagement was ensured through periodic comprehension questions (e.g., “Did you just see drinking?”) every 6-10 trials, requiring a yes or no button press.

### 2.5. Naive Experiment

This sequencing decision was influenced by pilot study results, which revealed that participants were able to accurately recognize the three agents (**Human**, **Android**, **Robot**) after watching the videos but struggled to do so with images alone. Consequently, the goal was to avoid any potential bias where insights acquired from the video sequences, especially regarding motion, could affect how participants interpreted the agents in static images. The **Naive Experiment** separated images and videos into two distinct blocks, with images always shown first, in order to prevent the acquisition of any potential prior knowledge about the agents from viewing the videos. Within each block, stimuli were pseudo-randomly sequenced to prevent backto-back repetitions. Every stimulus presentation began with a fixation cross for 900–1200 ms, randomized for timing. Comprehension questions similar to those in the **Prior Experiment** were asked intermittently to keep attention on the task, with responses given via key press.

### 2.6. EEG Data Collection and Preprocessing

Electroencephalography (EEG) was recorded using 64 ActiveTwo Ag/AgCl electrodes (manufactured by Brain Vision, Inc.) at a sampling rate of 512 Hz, adhering to the International 10/20 placement system. To ensure optimal signal quality, electrode impedances were maintained below 25 k-Ohm. Two additional electrodes were positioned above and below the right eye to track eye movements. EEG data was preprocessed using the EEGLAB software suite (Delorme and Makeig, 2004). Each participant’s data were subjected to high-pass filtering at 1 Hz and low-pass filtering at 50 Hz, which were then re-referenced to the average of the mastoid electrodes. In the **Naive Experiment**, visual examination revealed a noisy Fp2 electrode in two participants. Spherical interpolation was employed to correct for this noise in the affected subjects. After preprocessing, we segmented the data into epochs from 200 milliseconds before to 600 milliseconds after the video or first frame onset, time-locking these to the start of the video or static image to facilitate comparisons between **Moving** and **Still** agent representations. Epochs contaminated with eye movement or electromyographic artifacts were excluded from analysis using semi-automated rejection methods based on kurtosis and probability, with a standard deviation threshold set at 6.

### 2.7. Time-Resolved Representational Similarity Analysis

We utilized Time-Resolved Representational Similarity Analysis (RSA) (Kriegeskorte, 2008) to determine when the EEG signal begins to show distinguishable agent-specific signatures and to evaluate how these signals are altered by the absence of motion cues or prior knowledge. For this purpose, we generated 24 × 24 representational dissimilarity matrices (RDMs), encompassing all possible combinations of three agents and eight actions, at each EEG time point (refer to 2.7.1 for details). Additionally, we correlated these EEG-derived RDMs with a categorical Agent model (refer to 2.7.2 for details) using the Kendall-tau correlation coefficient. To further examine the influence of low-level visual features and other discernible effects, we developed two visual models (the *Pixel-wise Intensity Model* and the *Pixel-wise Motion Model*) alongside a categorical *Action Model*, as described in the 2.7.2 subsection.

#### 2.7.1. EEG Representational Dissimilarity Matrices

We generated EEG-based representational dissimilarity matrices (RDMs) for each participant and time point by computing the pairwise correlation distances across all channel data for each presented stimulus, resulting in a 24 × 24 matrix. Given the EEG’s sampling rate 512Hz, the interval between successive time points was 2 milliseconds. Recognizing that not all channels contribute equally to the signal, we further analyzed electrode subsets categorized by their scalp location. These subsets included **frontal** (Fz, F1, F2, F3, F4, AF3, AF4, AF7, AF8, AFz, Fp1, Fp2, FCz, FC1, FC2, FC5, FC6), **central** (Cz, C1, C2, C3, C4, C5, C6), **temporal** (T7, T8, FT9, FT10, FT7, FT8), **parietal** (Pz, P1, P2, P3, P4, P7, P8, POz, PO3, PO4, PO7, PO8), and **occipital** (Oz, O1, O2) electrodes. Using data from these specific groups of electrodes, we constructed region-specific EEG RDMs following the same methodology, employing custom Python scripts and functions for this task.

#### 2.7.2. Model Representational Dissimilarity Matrices

To dissect the temporal dynamics of agent-related information in EEG data and its modulation by different factors, we implemented a categorical attribute-based model, specifically the **Agent model**. However, significant correlations between the *Agent RDM* and *EEG RDMs* could be attributed to visual distinctions among stimuli (e.g., **Robot** stimuli may exhibit higher pixel intensities compared to **Human** stimuli). To account for this, we introduced three supplementary models: an attribute-based categorical *Action model*, along with a *Pixel-wise Motion model* and a *Pixel-wise Intensity model*. Subsequent analysis, detailed in subsection 2.7.3, aimed to identify when the **Agent model** significantly explains neural data. The attribute-based categorical model RDMs were constructed using either **Agent** attributes (*Robot*, *Human*, *Android*) or **Action** attributes (*Drink*, *Grasp*, *Handwave*, *Talk*, *Nudge*, *Throw*, *Turn*, *Wipe*), forming one-hot encoded vectors for each stimulus. In these vectors, a stimulus associated with a specific attribute was marked as 1 in the corresponding attribute entry and 0 elsewhere. We then calculated the pairwise Hamming distances between these binary vectors to generate a 24 × 24 RDM for each categorical model.

**Figure 1:**
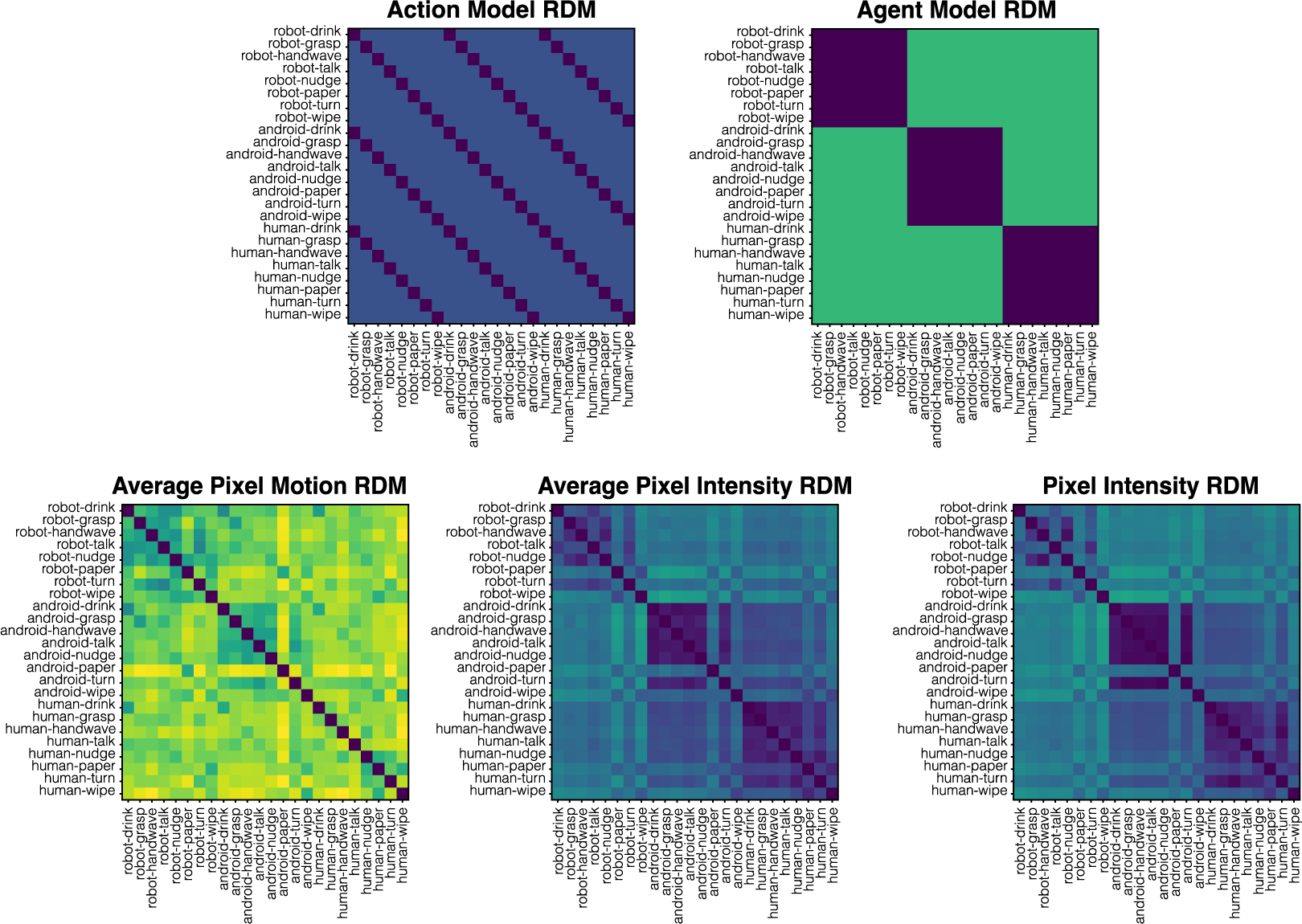
Model RDMs used in the multiple linear regression analysis.

For the low-level visual model RDMs, each stimulus was transformed into a feature vector based on the output of the relevant model (refer to the *Pixelwise Intensity RDM* and *Pixel-wise Motion RDM* sections for specifics) for that stimulus. Pairwise correlation distances between these feature vectors were calculated, producing a 24 × 24 RDM for each low-level model.

For the generation of low-level visual model RDMs, each stimulus was transformed into a vector reflecting the specific model’s output for that stimulus. We approached the pixel-wise intensity models with a distinction between **Moving** and **Still** conditions: for still images, we utilized the pixel intensities directly from the frames; for video stimuli, we averaged pixel intensities across all video frames. The pixel-wise motion model was constructed by calculating the optic flow for all pixels in each frame of the video stimuli and averaging these across frames (this model did not apply to the **Still** stimuli). These procedures yielded 2D matrices for pixel-wise intensity and motion for each stimulus, which were then converted into feature vectors. The correlation distance between each vector pair was computed, creating a 24 × 24 RDM for each low-level visual model.

#### 2.7.3. Multiple Linear Regression

We used time-resolved multiple linear regression analysis to examine the time course of specific EEG patterns that correspond to different agents. We controlled for potentially confounding factors like motion, pixel intensity, and action type. Our linear regression used the Model RDMs as independent variables to estimate the EEG RDM at each time point. This method helped us to identify the regression coefficients and their associated p-values for each model RDM at every time point. We also used Variance Inflation Factor (VIF) analysis to ensure that the model RDMs used in the multiple linear regression analysis did not have any multicollinearity issues.

## 3. Results

### 3.1. Whole-Brain RSA Results

Across the whole brain, representational similarity analysis revealed that peaks in agent identity information were consistently observed around 100 milliseconds post-stimulus in all conditions (*Naive-Moving*, *Naive-Still*, *Prior-Moving*, and *Prior-Still*). Distinctively, the *Naive Experiment* conditions uniquely reached a level of statistical significance, with p-values falling below 0.05 after adjustments for multiple comparisons via the Bonferroni correction, as illustrated in Figure 2. For the *Naive-Moving* condition, significant correlations between EEG RDMs and the Agent Model were found in the 118–144 ms interval. In contrast, the *Naive-Still* condition revealed a prolonged presence of agent information, with notable peaks at 70–118 ms and 154–196 ms intervals. Although the *Prior Experiment* yielded no significant findings after Bonferroni adjustment, the temporal profiles of *Prior-Still* and *Prior-Moving* closely mirrored those observed in the *Naive-Moving* condition.

**Figure 2:**
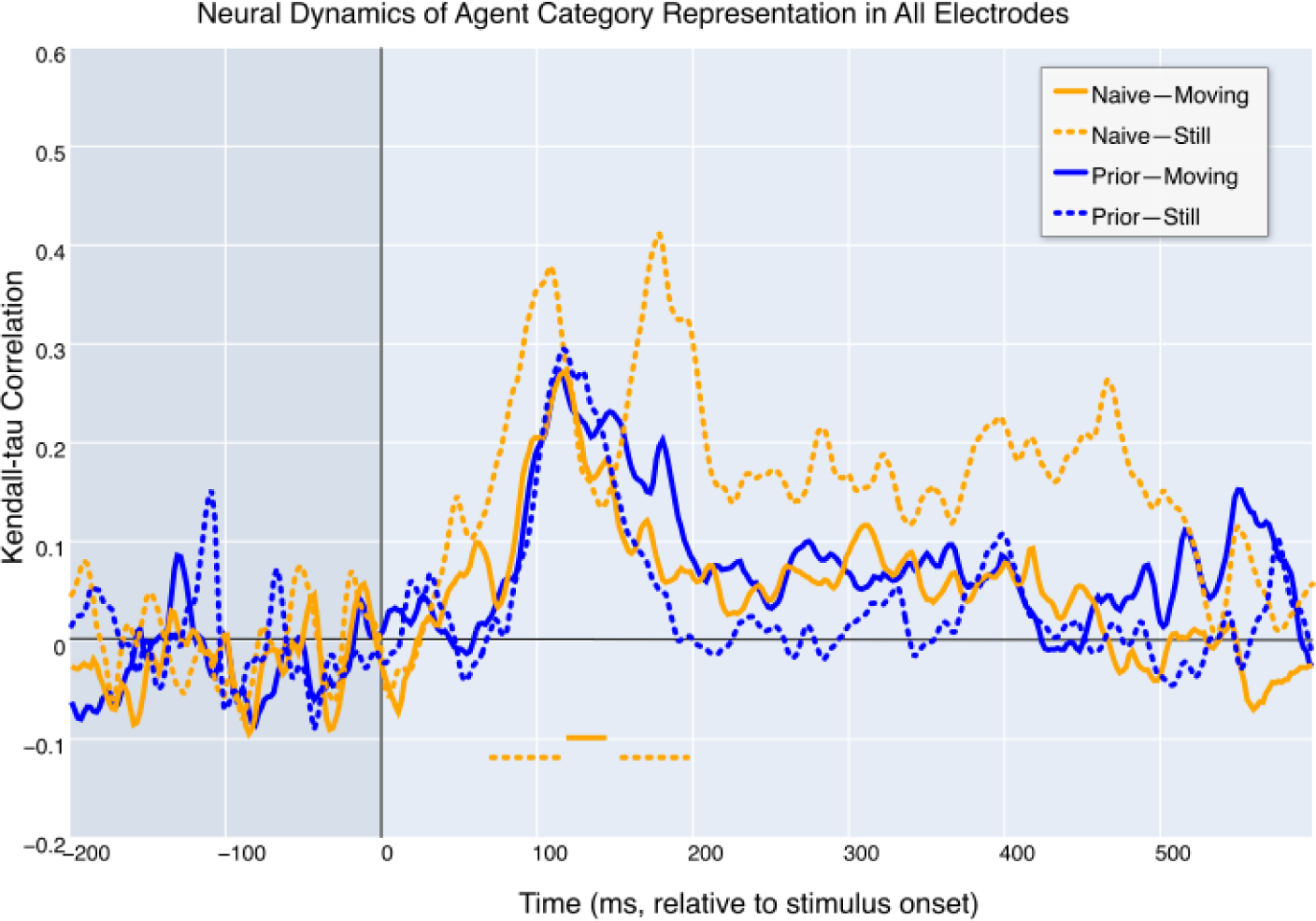
Temporal RSA across all electrodes. The figure maps time-resolved Kendall-tau correlations between EEG RDMs and the Agent RDM, corrected for confounding factors such as action identity, and form and motion properties at a low level. Time intervals where the correlations reach statistical significance after rigorous Bonferroni correction are indicated with horizontal lines below the correlation traces, with line styles matching the respective conditions.

### 3.2. Electrode-specific RSA

Our exploration into different sets of electrodes substantiated our preliminary predictions and exposed significant variability across electrode sites. While the central, frontal, and temporal electrode sites failed to demonstrate statistically significant correlations with the Agent model during any time segment, after accounting for the influences of alternative models, the occipital and parietal electrodes revealed intervals of statistical significance across all conditions, with p-values less than 0.05 (see Figure 3 for more details).

**Figure 3:**
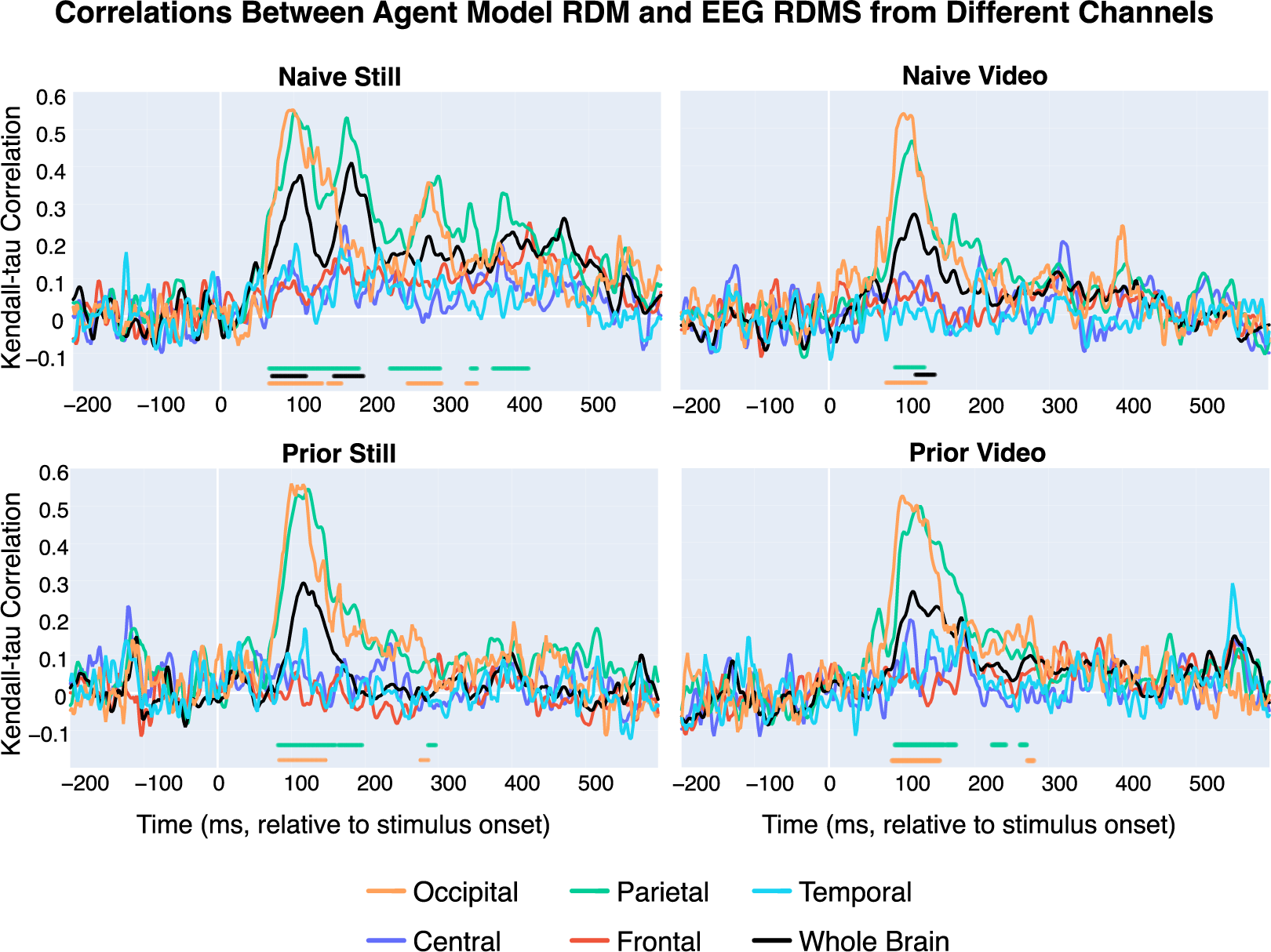
Temporal RSA across electrode sites. The figure maps time-resolved Kendall-tau correlations between EEG representational dissimilarity matrices (RDMs) and the Agent RDM, corrected for confounding factors such as action identity, and form and motion properties at a low level. Time intervals where the correlations reach statistical significance, after rigorous Bonferroni correction, are indicated with horizontal lines below the correlation traces, with line styles matching the respective electrode site.

The correlation patterns for both the occipital and parietal electrodes exhibit temporal profiles akin to those observed in the whole-brain correlations. However, the correlation magnitudes at the occipital and parietal sites are stronger than those across the entire set of electrodes, indicating a more reliable signal-to-noise ratio for this analysis. Presumably, the signals from occipital and parietal electrodes predominantly contributed to the effects initially observed in the whole-brain analysis (refer to Figure 2). When the analysis is confined to just the occipital or parietal electrodes, we identify statistically significant time periods for the **Prior Experiment**, which were not evident when the entire electrode array was considered. Therefore, subsequent investigations will be tailored to these most informative electrode groups, specifically the occipital and parietal electrodes.

### 3.3. RSA on Parietal and Occipital Electrodes

Upon controlling for the confounding effects of action identity and the elementary visual factors of motion and pixel intensity, a significant correlation was observed between the EEG responses at the parietal electrodes and the Agent model. This correlation was evident from 66 to 418 milliseconds in the *Naive-Still* scenario, from 90 to 130 milliseconds in the **Naive-Moving** scenario, from 82 to 296 milliseconds in the **Prior-Still** scenario, and from 90 to 242 milliseconds in the **Prior-Moving** scenario, as depicted in Figure 4.

**Figure 4:**
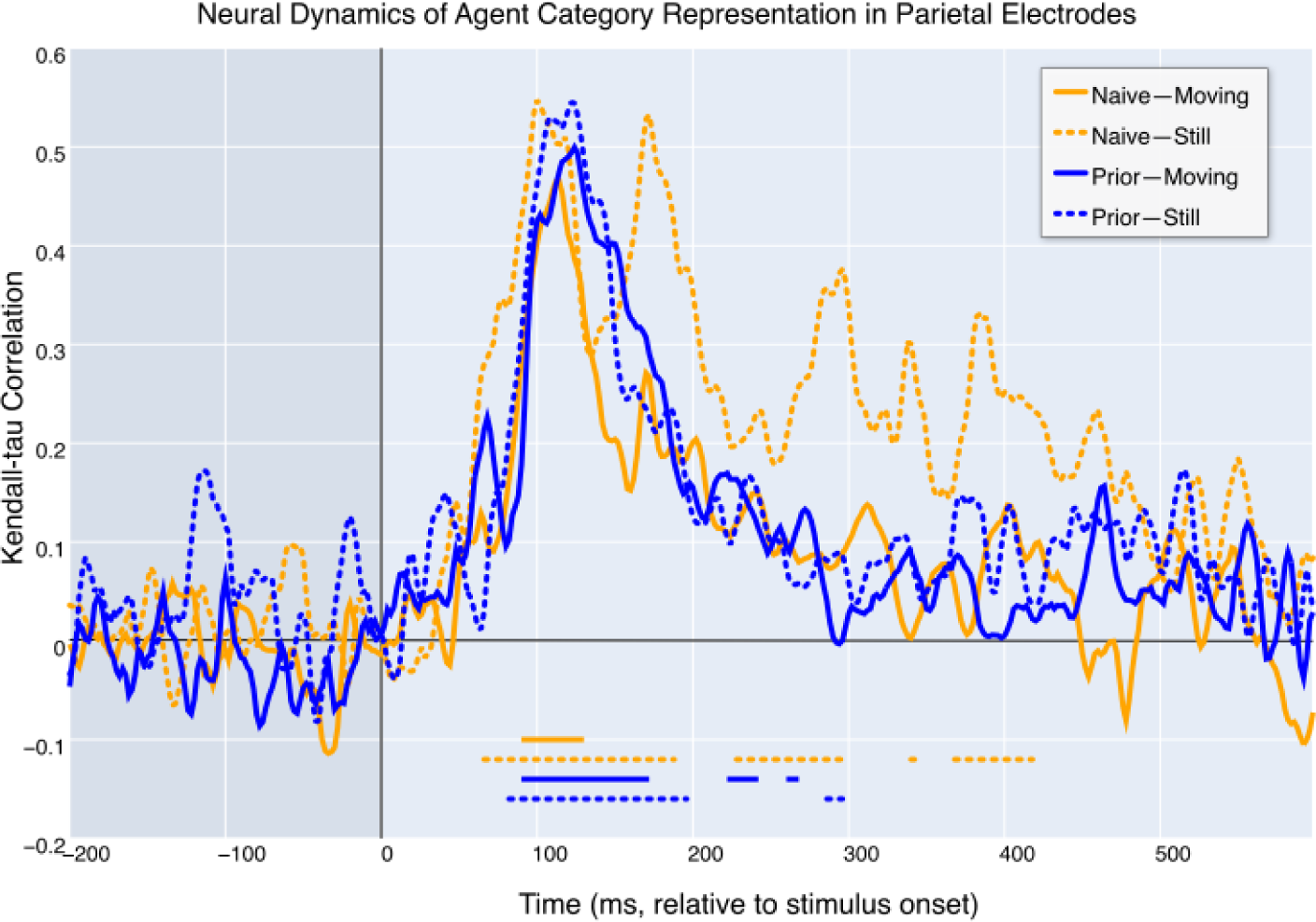
Temporal RSA across parietal electrodes. The figure maps time-resolved Kendall-tau correlations between EEG RDMs and the Agent RDM, corrected for confounding factors such as action identity, and form and motion properties at a low level. Time intervals where the correlations reach statistical significance after Bonferroni correction are indicated with horizontal lines below the correlation traces, with line styles matching the respective conditions.

Results from the occipital electrode analysis were consistent with those of the parietal electrodes, as evidenced in Figure 3. Notably, parietal electrodes exhibited higher peak values in comparison to occipital electrodes across all conditions. Specifically, in the **Naive-Still** condition, there was a distinctive persistence of correlation beyond the initial peak. Parietal electrodes more effectively highlighted later peaks in correlation; therefore, for the sake of visual clarity, we have chosen to present comparisons of conditions exclusively for the parietal electrodes.

For the **Prior-Moving**, **Prior-Still**, and **Naive-Moving** conditions, the correlation patterns are strikingly similar: an ascent precedes the 100 ms mark, culminating in a peak around 120 ms, followed by a slow descent approaching 200 ms. Beyond 200 ms, a subtle effect emerges, sometimes achieving significance between 200 and 300 ms in the **Prior Experiment**. Despite the **Naive-Moving** condition not attaining statistical significance after its initial peak, its trajectory remains consistent with the patterns delineated for the **Prior Experiment**.

The **Naive-Still** condition stands out from the rest, characterized by a correlation trace with several peaks and achieving statistical significance during considerably later phases. Unlike the other conditions, the **Naive-Still** condition’s correlation trace spans a wider interval, notably from 150 to 450 ms. Importantly, the peak occurring around 400 ms is distinguished by its statistical significance.

## 4. Discussion

The skill to comprehend actions and the agents executing them is central to human cognition. Nevertheless, the temporal patterns of agent perception and the interplay between bottom-up and top-down mechanisms demand a more in-depth investigation.

Our study investigated the interaction between prior knowledge and motion information on the temporal processing of agents varying in human likeness. We manipulated motion information (**Still** vs. **Moving**) and prior knowledge (**Prior** vs. **Naive**), and employed scalp EEG with model-based temporal representational similarity analysis (RSA). We posited that motion cues and prior knowledge would impact the temporal signature of agent representation, especially under conditions where motion cues are absent.

Our results suggest that when there is no prior knowledge about the agents and no motion cues in the stimuli, the correlation between the agent RDM and EEG RDMs has a longer duration than in other conditions. This observation can be attributed to several factors: First, motion cues are pivotal for the recognition of biological motion and the observation of actions, as suggested by prior research (Giese and Poggio, 2003; Blake and Shiffrar, 2007; Garcia and Grossman, 2008). Given the close relationship between agent perception and biological motion, this also suggests that the absence of motion cues likely necessitated prolonged processing of agent information. Second, the continued representation observed in the EEG signal might reflect a kind of recurrent process. This recurrent process may be needed to clarify ambiguity when there are no motion cues or prior knowledge. This suggests that a more concise representation could indicate less dependence on such recurrent processing. Third, without motion cues participants might depend more on other visual cues, such as the agents’ shape, texture, or other features. This reliance could enhance the influence of agent representations on EEG signals. Although we attempted to mitigate the impact of low-level visual details on our correlations through regression with low-level visual models, these models may not fully account for all visual differences, like texture or shape, that participants might utilize more in conditions devoid of motion cues and prior knowledge.

### 4.1. Prior information eliminates the advantage of motion information

Our findings highlight that the lack of motion cues extends the duration of significant correlations between the Agent RDM and EEG RDMs, but this effect is exclusively observed in the **Naive Experiment**. In this experiment, participants lacked prior information about the agents and their actions. Conversely, in the **Prior Experiment**, the significance of correlation patterns across both **Still** and **Moving** conditions was comparable, implying that prior knowledge about the agents and their actions diminishes the need for motion information.

The reduced need for motion cues, when prior knowledge is available, may have various causes. in the **Naive-Still** condition, without both prior knowledge and motion cues, distinguishing agent identities becomes more complex, thereby prolonging and intensifying the correlation. This observation is consistent with Bayesian inference models, which postulate that ambiguous stimuli necessitate extended and intensified processing due to the prediction errors they cause, prompting recurrent updates (Kaiser et al., 2023; Urgen and Saygin, 2020; Summerfield and de Lange, 2014; Kersten et al., 2004).

Alternatively, prior knowledge about the agents may reduce prediction errors, thereby diminishing neural activation (Press et al., 2020; Summerfield and de Lange, 2014). It is also conceivable that such knowledge enables topdown modulation of perception, helping process essential agent and action features using the remaining visual cues in the **Still** condition. Future investigations should adopt more refined controls for low-level visual influences. This necessitates sophisticated strategies that go beyond regression against low-level visual models. Studies by Gosselin and Schyns (2001); Thurman and Grossman (2008) offer insights into methodologies that could achieve this nuanced control.

### 4.2. Ambiguity around Agent Identities can be Efficiently Resolved by either Motion Cues or Prior Information

In this study, direct influences from either motion availability or prior knowledge on agent representation over time were not observed in the overall temporal RSA results.

The impact of motion became apparent only in the **Naive Experiment** and was specific to the differences between the **Naive-Still** and **Naive-Moving** conditions. Similarly, the influence of prior knowledge was observed only within the **Moving** conditions, particularly when comparing the **Prior-Moving** and **Naive-Moving** conditions. Interestingly, a unique temporal pattern of agent identity representation emerged exclusively in the **Naive-Still** condition. In contrast, the remaining conditions (**Prior-Still**, **Prior-Moving**, and **Naive-Moving**) displayed similar temporal representation profiles. This indicates that the presence of either motion cues or prior knowledge alone effectively reduces the ambiguity associated with agent identities. Shorter agent representation in EEG may indicate a more efficient resolution of ambiguity around agent identities in the presence of motion cues or prior knowledge. This suggests that, in our case, incorporating either motion cues or prior knowledge is enough to facilitate agent processing, leading to a faster resolution of ambiguity in agent identities. Future research is necessary to understand further the connection between motion, form, and prior information in situations where ambiguity is higher, for instance, due to additional sensory noise or the introduction of parametrical manipulation of form and motion cues. Based on our findings, we anticipate that the reliance on prior information should increase as the level of ambiguity in form and motion information gradually increases. This would also lead to a gradual change in how much motion and form cues can resolve the ambiguity in the agent identities and, therefore, change the duration of agent representation. The prolonged representation of agents observed in the **Naive-Still** condition may be due to increased recurrent processing, which is potentially necessary to disambiguate identities in the absence of motion cues and prior knowledge (Kietzmann et al., 2019). This suggests that a more concise representation may indicate reduced reliance on such recurrent processing for disambiguation. Future studies utilizing recurrent models can investigate the relationship between the dependence on recurrent processing, the ambiguity in agent identities, and their relationship with the amount of low-level sensory information.

## 5. Conclusion

In conclusion, our research explored the temporal dynamics of agent processing, focusing on the effects of prior knowledge and motion cues. Employing time-resolved representational similarity analysis and regression techniques, we discovered that the presence of motion and prior knowledge significantly influence agent processing in EEG recordings. Notably, in the naive condition, agent information persisted for longer periods during still presentations than moving ones. Conversely, the duration of agent information processing was similar across still and moving scenarios in the prior knowledge condition. These observations indicate that prior knowledge and motion cues interact in a way that impacts the time agent information is processed. Our findings emphasize the importance of both prior knowledge and motion cues in the cognitive processing of agent information. They showcase the intricate relationship between top-down modulation and bottom-up signals in the perception of agents and their actions.

## Acknowledgements

The authors would like to thank Ayse Pinar Saygin for her support during the data acquisition of the project.

## 7. CRediT authorship contribution statement

Sena Er: Methodology, Software, Formal Analysis, Visualization, Writing – original draft, Writing – review & editing. Huseyin O. Elmas: Methodology, Software, Formal Analysis, Visualization, Writing – original draft, Writing – review & editing. Burcu A. Urgen: Conceptualization, Investigation, Funding acquisition, Resources, Supervision, Writing – review & editing.

## References

Bach, P., Schenke, K.C., 2017. Predictive social perception: Towards a unifying framework from action observation to person knowledge. Social and Personality Psychology Compass 11, e12312. doi:10/gg6xdq.

Bar, M., Kassam, K.S., Ghuman, A.S., Boshyan, J., Schmid, A.M., Dale, A.M., Hämäläinen, M.S., Marinkovic, K., Schacter, D.L., Rosen, B.R., Halgren, E., 2006. Top-down facilitation of visual recognition. Proceedings of the National Academy of Sciences 103, 449–454. doi:10/cxh5xr.

Beintema, J.A., Lappe, M., 2002. Perception of biological motion without local image motion. Proceedings of the National Academy of Sciences 99, 5661–5663. doi:10/fqf9zj.

Bill, J., Gershman, S.J., Drugowitsch, J., 2022. Visual motion perception as online hierarchical inference. Nature Communications 13, 7403. doi:10/gtn7m5.

Blake, R., Shiffrar, M., 2007. Perception of Human Motion. Annual Review of Psychology 58, 47–73. doi:10/fr7p9h.

Caspers, S., Zilles, K., Laird, A.R., Eickhoff, S.B., 2010. ALE meta-analysis of action observation and imitation in the human brain. NeuroImage 50, 1148–1167. doi:10/dhxprp.

Chambon, V., Domenech, P., Jacquet, P.O., Barbalat, G., Bouton, S., Pacherie, E., Koechlin, E., Farrer, C., 2017. Neural coding of prior expectations in hierarchical intention inference. Scientific Reports 7, 1–16. doi:10/f96qn3.

Cross, E.S., Liepelt, R., de C. Hamilton, A.F., Parkinson, J., Ramsey, R., Stadler, W., Prinz, W., 2011. Robotic movement preferentially engages the action observation network. Human Brain Mapping 33, 2238–2254. doi:10/b6dzn4.

Delorme, A., Makeig, S., 2004. EEGLAB: An open source toolbox for analysis of single-trial EEG dynamics including independent component analysis. Journal of Neuroscience Methods 134, 9–21. doi:10/bqr2f2.

Flounders, M.W., González-García, C., Hardstone, R., He, B.J., 2019. Neural dynamics of visual ambiguity resolution by perceptual prior. eLife 8, e41861. doi:10/gfw7gm.

Garcia, J.O., Grossman, E.D., 2008. Necessary but not sufficient: Motion perception is required for perceiving biological motion. Vision Research 48, 1144–1149. doi:10/bftrxw.

Gazzola, V., Rizzolatti, G., Wicker, B., Keysers, C., 2007. The anthropomorphic brain: The mirror neuron system responds to human and robotic actions. NeuroImage 35, 1674–1684. doi:10/cv3xmf.

Giese, M.A., Poggio, T., 2003. Neural mechanisms for the recognition of biological movements. Nature Reviews Neuroscience 4, 179–192. doi:10/c25gzb.

Gilaie-Dotan, S., Saygin, A.P., Lorenzi, L.J., Egan, R., Rees, G., Behrmann, M., 2013. The role of human ventral visual cortex in motion perception. Brain 136, 2784–2798. doi:10/f48ks9.

Gosselin, F., Schyns, P.G., 2001. Bubbles: A technique to reveal the use of information in recognition tasks. Vision Research 41, 2261–2271. doi:10/cdx752.

Hardstone, R., Zhu, M., Flinker, A., Melloni, L., Devore, S., Friedman, D., Dugan, P., Doyle, W.K., Devinsky, O., He, B.J., 2021. Long-term priors influence visual perception through recruitment of long-range feedback. Nature Communications 12, 1–15. doi:10/gtn7m3.

Hunt, A.R., Halper, F., 2008. Disorganizing biological motion. Journal of Vision 8, 12–12. doi:10/fbz4zh.

Jacquet, P.O., Roy, A.C., Chambon, V., Borghi, A.M., Salemme, R., Farné, A., Reilly, K.T., 2016. Changing ideas about others’ intentions: Updating prior expectations tunes activity in the human motor system. Scientific Reports 6, 1–15. doi:10/f8qng5.

Jellema, T., Perrett, D.I., 2003. Cells in monkey STS responsive to articulated body motions and consequent static posture: A case of implied motion? Neuropsychologia 41, 1728–1737. doi:10/crv5qr.

Kaiser, D., Stecher, R., Doerschner, K., 2023. EEG Decoding Reveals Neural Predictions for Naturalistic Material Behaviors. The Journal of Neuroscience 43, 5406–5413. doi:10/gtn7nd.

Kar, K., Kubilius, J., Schmidt, K., Issa, E.B., DiCarlo, J.J., 2019. Evidence that recurrent circuits are critical to the ventral stream’s execution of core object recognition behavior. Nature Neuroscience 22, 974–983. doi:10/gfz7dv.

Kersten, D., Mamassian, P., Yuille, A., 2004. Object Perception as Bayesian Inference. Annual Review of Psychology 55, 271–304. doi:10/djzsrk.

Kietzmann, T.C., Spoerer, C.J., Sörensen, L.K.A., Cichy, R.M., Hauk, O., Kriegeskorte, N., 2019. Recurrence is required to capture the representational dynamics of the human visual system. Proceedings of the National Academy of Sciences 116, 21854–21863. doi:10/gf9j2t.

Kornmeier, J., Bhatia, K., Joos, E., 2021. Top-down resolution of visual ambiguity – knowledge from the future or footprints from the past? PLOS ONE 16, e0258667. doi:10/grd6tg.

Kriegeskorte, N., 2008. Representational similarity analysis – connecting the branches of systems neuroscience. Frontiers in Systems Neuroscience 2, 1–28. doi:10/fh5dft.

Kupferberg, A., Iacoboni, M., Flanagin, V., Huber, M., Kasparbauer, A., Baumgartner, T., Hasler, G., Schmidt, F., Borst, C., Glasauer, S., 2017. Fronto-parietal coding of goal-directed actions performed by artificial agents. Human Brain Mapping 39, 1145–1162. doi:10/gcnsfp.

Lange, J., Lappe, M., 2006. A Model of Biological Motion Perception from Configural Form Cues. The Journal of Neuroscience 26, 2894–2906. doi:10/fj3ch8.

Markov, N.T., Vezoli, J., Chameau, P., Falchier, A., Quilodran, R., Huissoud, C., Lamy, C., Misery, P., Giroud, P., Ullman, S., Barone, P., Dehay, C., Knoblauch, K., Kennedy, H., 2013. Anatomy of hierarchy: Feedforward and feedback pathways in macaque visual cortex. Journal of Comparative Neurology 522, 225–259. doi:10/gfzd9x.

McLeod, P., 1996. Preserved and Impaired Detection of Structure From Motion by a ‘Motion-blind” Patient. Visual Cognition 3, 363–392. doi:10/cvxmd5.

Mishkin, M., Ungerleider, L.G., Macko, K.A., 1983. Object vision and spatial vision: Two cortical pathways. Trends in Neurosciences 6, 414–417. doi:10/c4kbpn.

Nelissen, K., Borra, E., Gerbella, M., Rozzi, S., Luppino, G., Vanduffel, W., Rizzolatti, G., Orban, G.A., 2011. Action Observation Circuits in the Macaque Monkey Cortex. The Journal of Neuroscience 31, 3743–3756. doi:10/cpkms9, arXiv:https://www.jneurosci.org/content/31/10/3743.full.pdf.

Neri, P., Morrone, M.C., Burr, D.C., 1998. Seeing biological motion. Nature 395, 894–896. doi:10/fjxvjr.

Ondobaka, S., de Lange, F.P., Wittmann, M., Frith, C.D., Bekkering, H., 2014. Interplay Between Conceptual Expectations and Movement Predictions Underlies Action Understanding. Cerebral Cortex 25, 2566–2573. doi:10/f7rgf6.

Orban, G.A., 2018. Action Observation as a Visual Process: Different Classes of Actions Engage Distinct Regions of Human PPC. Exploring Complexity, 1–32 doi:10/gtn7mz.

O’Reilly, R.C., Wyatte, D., Herd, S., Mingus, B., Jilk, D.J., 2013. Recurrent Processing during Object Recognition. Frontiers in Psychology 4, 124. doi:10/gc7rk6.

Pantle, A., Turano, K., 1992. Visual resolution of motion ambiguity with periodic luminanceand contrast-domain stimuli. Vision Research 32, 2093– 2106. doi:10/cntxsb.

Pitcher, D., Ungerleider, L.G., 2021. Evidence for a Third Visual Pathway Specialized for Social Perception. Trends in Cognitive Sciences 25, 100–110. doi:10/ghpvc5.

Platonov, A., Orban, G.A., 2016. Action observation: The less-explored part of higher-order vision. Scientific Reports 6, 1–13. doi:10/f9cfg4.

Press, C., Kok, P., Yon, D., 2020. The Perceptual Prediction Paradox. Trends in Cognitive Sciences 24, 13–24. doi:10/ggd6zp.

Salge, J.H., Pollmann, S., Reeder, R.R., 2020. Anomalous visual experience is linked to perceptual uncertainty and visual imagery vividness. Psychological Research 85, 1848–1865. doi:10/gtn7m2.

Saygin, A.P., Chaminade, T., Ishiguro, H., Driver, J., Frith, C., 2011. The thing that should not be: Predictive coding and the uncanny valley in perceiving human and humanoid robot actions. Social Cognitive and Affective Neuroscience 7, 413–422. doi:10/cmx78w.

Schenke, K.C., Wyer, N.A., Bach, P., 2016. The Things You Do: Internal Models of Others’ Expected Behaviour Guide Action Observation. PLOS ONE 11, e0158910. doi:10/gbpmvp.

Sliwa, J., Freiwald, W.A., 2017. A dedicated network for social interaction processing in the primate brain. Science 356, 745–749. doi:10/f98mhd.

Sterzer, P., Kleinschmidt, A., Rees, G., 2009. The neural bases of multistable perception. Trends in Cognitive Sciences 13, 310–318. doi:10/dqgptr.

Summerfield, C., de Lange, F.P., 2014. Expectation in perceptual decision making: Neural and computational mechanisms. Nature Reviews Neuroscience 15, 745–756. doi:10/gf9qhk.

Tarhan, L., Konkle, T., 2020. Sociality and interaction envelope organize visual action representations. Nature Communications 11, 1–11. doi:10/gqvjwd.

Thurman, S.M., Giese, M.A., Grossman, E.D., 2010. Perceptual and computational analysis of critical features for biological motion. Journal of Vision 10, 15–15. doi:10/bxnq8r.

Thurman, S.M., Grossman, E.D., 2008. Temporal “Bubbles” reveal key features for point-light biological motion perception. Journal of Vision 8, 28. doi:10/bbpq85.

Urgen, B.A., Kutas, M., Saygin, A.P., 2018. Uncanny valley as a window into predictive processing in the social brain. Neuropsychologia 114, 181–185. doi:10/gd9drq.

Urgen, B.A., Saygin, A.P., 2020. Predictive processing account of action perception: Evidence from effective connectivity in the action observation network. Cortex 128, 132–142. doi:10/gpfk5d.

Vaina, L.M., Lemay, M., Bienfang, D.C., Choi, A.Y., Nakayama, K., 1990. Intact “biological motion” and “structure from motion” perception in a patient with impaired motion mechanisms: A case study. Visual Neuroscience 5, 353–369. doi:10/dbc5d4.

van Bergen, R.S., Kriegeskorte, N., 2020. Going in circles is the way forward: The role of recurrence in visual inference. Current Opinion in Neurobiology 65, 176–193. doi:10/gqvjsz.

Wurm, M.F., Caramazza, A., Lingnau, A., 2016. Action Categories in Lateral Occipitotemporal Cortex Are Organized Along Sociality and Transitivity. The Journal of Neuroscience 37, 562–575. doi:10/gtn7mx.

Wyatte, D., Jilk, D.J., O’Reilly, R.C., 2014. Early recurrent feedback facilitates visual object recognition under challenging conditions. Frontiers in Psychology 5. doi:10/f59p4j.

